# Traffickome infers directed, compartment-resolved membrane trafficking from proteomic data

**DOI:** 10.64898/2026.09.20.753016

**Authors:** Rushikesh Deshpande, Itziar Pinilla-Macua, Joao Paulo, Steven P. Gygi, Alexander Sorkin

## Abstract

Membrane trafficking determines the subcellular localization and compartment-residence time of receptors, channels, transporters and other transmembrane proteins and thereby influences where in the cell and how long these proteins function. Functional-enrichment computational approaches may predict the membrane-trafficking regulations but do not resolve the directionality of the transport between cellular compartments. We present Traffickome, a deterministic method organizing 754 trafficking proteins into 11 directed transitions across seven compartments and 1,837 signaling proteins across 22 pathways. Implemented as an interactive browser and open-source Python package, Traffickome infers compartment-to-compartment transport, cargo fate and signaling-pathway activity and enables in silico perturbation from a single proteomic comparison. Across 29 proteomic comparisons spanning 14 studies encompassing over 66,000 gene-level observations across over 10,000 genes, Traffickome ranked the expected trafficking transition first in 23 cases (79%), outperforming the best conventional enrichment methods (17 of 29, 59%). Mean accuracy fell to 18% after randomizing the protein-to-transition assignments. To illustrate the power of Traffickome, we performed time-resolved analysis of the phosphoproteome in EGF-stimulated cells with the goal to distinguish signaling by internalized versus plasma-membrane EGFR. In the resulting datasets, comprising over 18,000 phosphosite-level measurements, Traffickome revealed localization-dependent differences in EGF-induced phosphorylation and prioritized trafficking regulators for testing their effects on signaling processes.

## Introduction

Membrane trafficking governs the intracellular movement of transmembrane cargoes – receptors, channels, transporters, and resident membrane proteins [1]. For signaling receptors, this movement determines location and duration of receptor-associated signaling [1]. The same ligand-bound receptor can trigger diverse signaling outcomes depending on whether it stays at the plasma membrane, localizes at endosomes, or is delivered to lysosomes [1]. Intracellular trafficking is commonly dysregulated in diseases such as cancer and neurodegeneration [2,3]. Therefore, approaches that can infer directional trafficking from proteomic measurements are valuable.

Microscopy and biochemical trafficking assays remain the standard methods for tracking a single cargo between intracellular compartments. Live-cell imaging and pulse-chase experiments track labeled cargo as it moves from one compartment to another. Multiplexed imaging extends these observations to additional organelle markers. However, these methods require engineered or labeled cargo and are restricted to proteins selected in advance. Mass spectrometry routinely quantifies thousands of proteins in a single experiment, including many trafficking regulators and signaling effectors. The enrichment and pathway-analysis methods commonly applied to these datasets treat trafficking proteins as unordered gene sets [4–8].

They identify enrichment of proteins assigned to particular cellular compartments or trafficking processes but do not determine whether transport is directed towards or away from those compartments. Several computational methods infer protein relocalization from spatial proteomics. pRoloc, BANDLE, and TRANSPIRE identify changes in protein localization between subcellular compartments [9–11]. These methods require spatially resolved proteomic measurements and describe the localization or relocalization of individual proteins without scoring directed compartment-to-compartment trafficking transitions. Directional membrane trafficking is therefore not addressed by the standard analyses of differential and proximity-proteomic datasets.

We present Traffickome, a deterministic method that infers directional, compartment-resolved membrane trafficking from a single proteomic comparison without requiring spatial fractionation or model training. Traffickome converts a curated directional trafficking reference set into a deterministic scoring algorithm to quantify support for transport between specific compartments given any proteomic dataset. A parallel signaling reference set enables interpreting transition and signaling-pathway scores together. In this method, cargo denotes a membrane protein being transported, including receptors, channels, transporters, and resident membrane proteins, while receptor-associated cytosolic proteins are analyzed separately as signaling mediators.

Traffickome projects trafficking transition scores onto cargo fates for membrane cargoes, implements in silico perturbation of trafficking regulators, and compares proteomic predictions with evidence for protein localizations extracted from the literature using fluorescence microscopy. The method is available as an interactive web application and as an open-source Python package.

## Results

### Traffickome organizes proteomic data into directed trafficking transitions

Traffickome describes membrane trafficking by 11 directed transitions connecting seven compartments across endocytic, recycling, degradative, secretory and retrograde transport (Fig. 1a). The trafficking reference set includes 754 proteins, and the signaling reference set includes 1,837 proteins mapped to 22 pathways (Fig. 1b). The same protein can appear in multiple transitions or pathways (Fig. 1c,d). Each proteomics dataset is converted into gene-level scores and classified as supporting each transition specifically, broadly or not at all. This profile is used for cargo-fate projection, in silico perturbation and comparison with fluorescence-imaging evidence.

**Figure 1.**
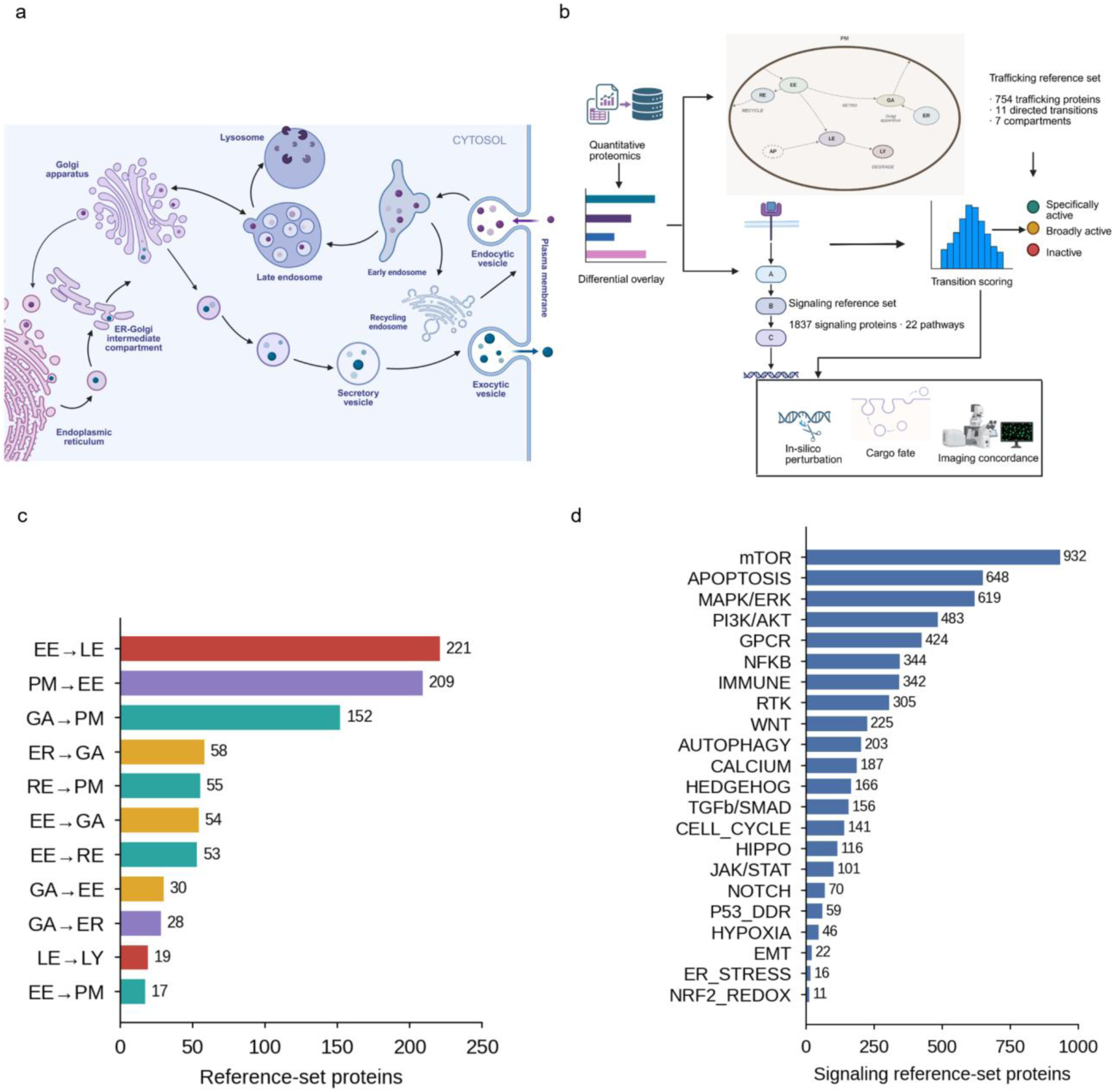
Traffickome framework and reference sets. **a.** Schematic of the seven route-endpoint compartments and the eleven directed transitions connecting them. **b.** Analysis workflow, from proteomic input through gene-score standardization, transition scoring and permutation testing to cargo-fate projection and in silico perturbation. **c.** Number of protein-transition assignments for each of the eleven directed transitions, coloured by cargo-fate class; 896 assignments across 752 proteins. **d.** Number of proteins assigned to each of the 22 signaling pathways; 1,837 proteins total, with proteins assigned to more than one pathway counted once per pathway. PM, plasma membrane; EE, early endosome; LE, late endosome; LY, lysosome; RE, recycling endosome; GA, Golgi apparatus; ER, endoplasmic reticulum. Panel a was created with BioRender.com.

### The Traffickome browser connects transition inference to mechanistic insight

Traffickome is available as a browser application in which uploaded data are processed locally. As an example, using the 30 min timepoint from a published time-resolved EGFR-APEX2 proximity-labeling time course in HCT116 cells [12], the Upload tab identifies the analysis mode and filtering parameters (Fig. 2a), and the Transitions tab ranks directed transport evidence, with EE→LE as the highest-ranked specific transition (Fig. 2b). HGS, a hepatocyte growth factor-regulated tyrosine kinase substrate (alternative common protein name is HRS) [13], was selected as an illustrative perturbation target. HGS is an ESCRT-0 subunit that recognizes ubiquitinated cargo at early endosomes and mediates sorting of such cargo for the lysosomal degradation pathway [13]. Simulated HGS knockout with network propagation produced its largest decrease in EE→LE support (ΔPₑ = −1.157; Fig. 2c), linking directional inference to mechanistic output.

**Figure 2.**
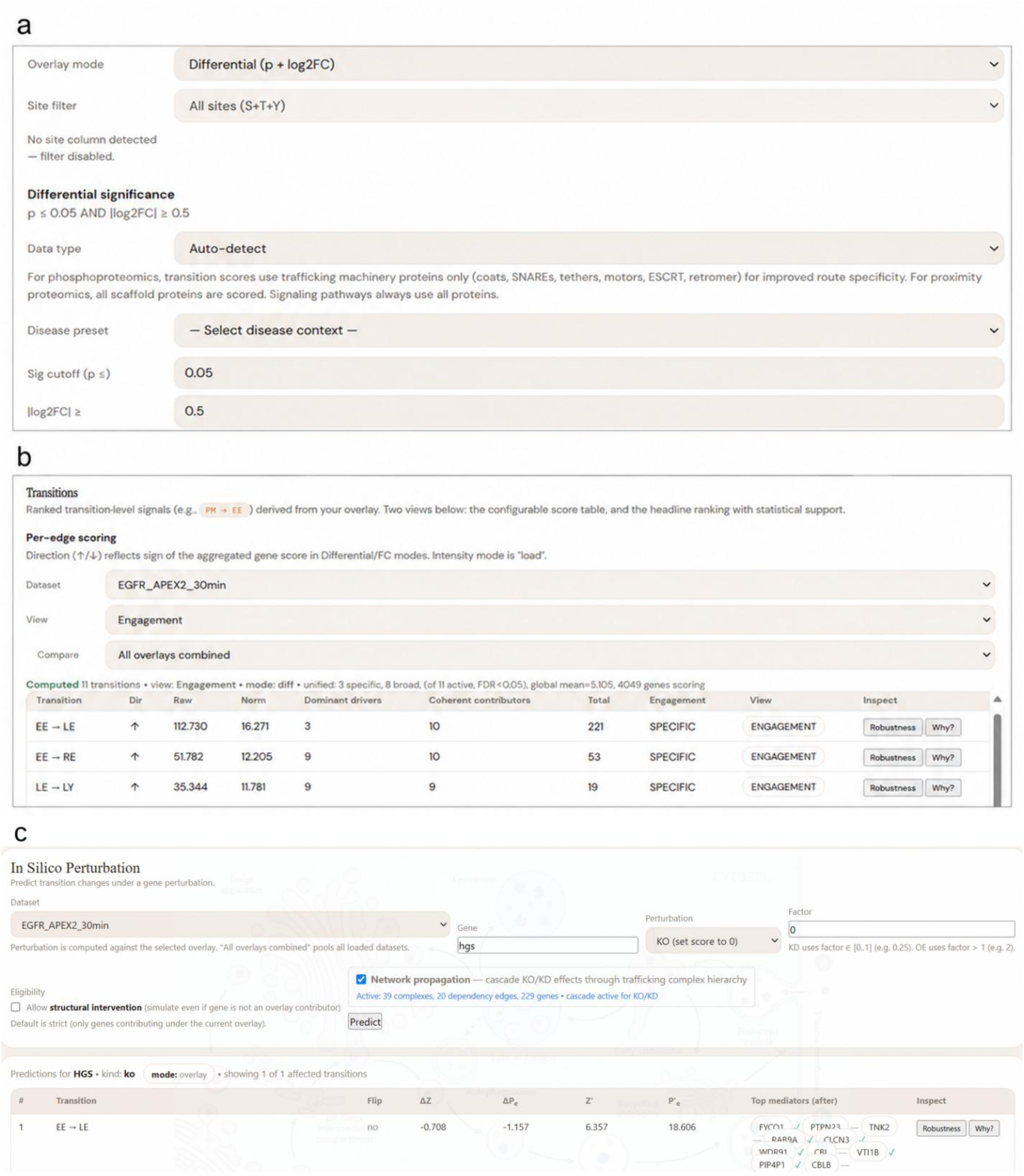
Browser workflow for directional trafficking analysis. Screenshots of the browser application analysing a published 30-min EGFR-APEX2 proximity-labelling dataset. **a.** Import settings, showing automatic detection of the differential input mode and the default gates (P ≤ 0.05, |log2 fold change| ≥ 0.5). **b.** Ranked transition output. Transitions are ordered by absolute adjusted score; EE→LE is the leading SPECIFIC transition (Pₑ = 16.271) across 4,049 scoring genes. **c.** In silico HGS knockout with complex-dependency propagation enabled. EE→LE is the only affected transition, decreasing by ΔPₑ = −1.157 (ΔZ = −0.708), with the surviving contributors listed. All scoring is performed locally in the browser; uploaded data are not transmitted.

### Traffickome infers cargo trafficking fate from proteomic data

We first used Traffickome to infer the endocytic progression of EGFR from a published APEX2 proximity-labeling time course collected at 1, 10, 30 and 60 min after EGF stimulation [12]. Scoring against the 754-protein directional trafficking reference set identified the expected temporal sequence of compartment-to-compartment transition support (Fig. 3a). At 1 min, EGFR remained predominantly plasma-membrane associated and the cargo projection favored plasma-membrane retention. The PM→EE score likely reflects the recruitment of clathrin-mediated endocytic machinery rather than movement into early endosomes. By 10 min, ligand-bound EGFR was largely in the early/sorting endosomes and the cargo projection favored further movement into this compartment. This transition support became significant bringing the EE→LE score to its specific maximum. The cargo projection favored movement directly into the degradative machinery (four transition supports) rather than ER. At 30 and 60 min, late-endosomal and recycling transition support followed (Fig. 3a). This progression was consistent with the established movement of ligand-activated EGFR from the plasma membrane through the endosomal system towards lysosomal degradation [1].

**Figure 3.**
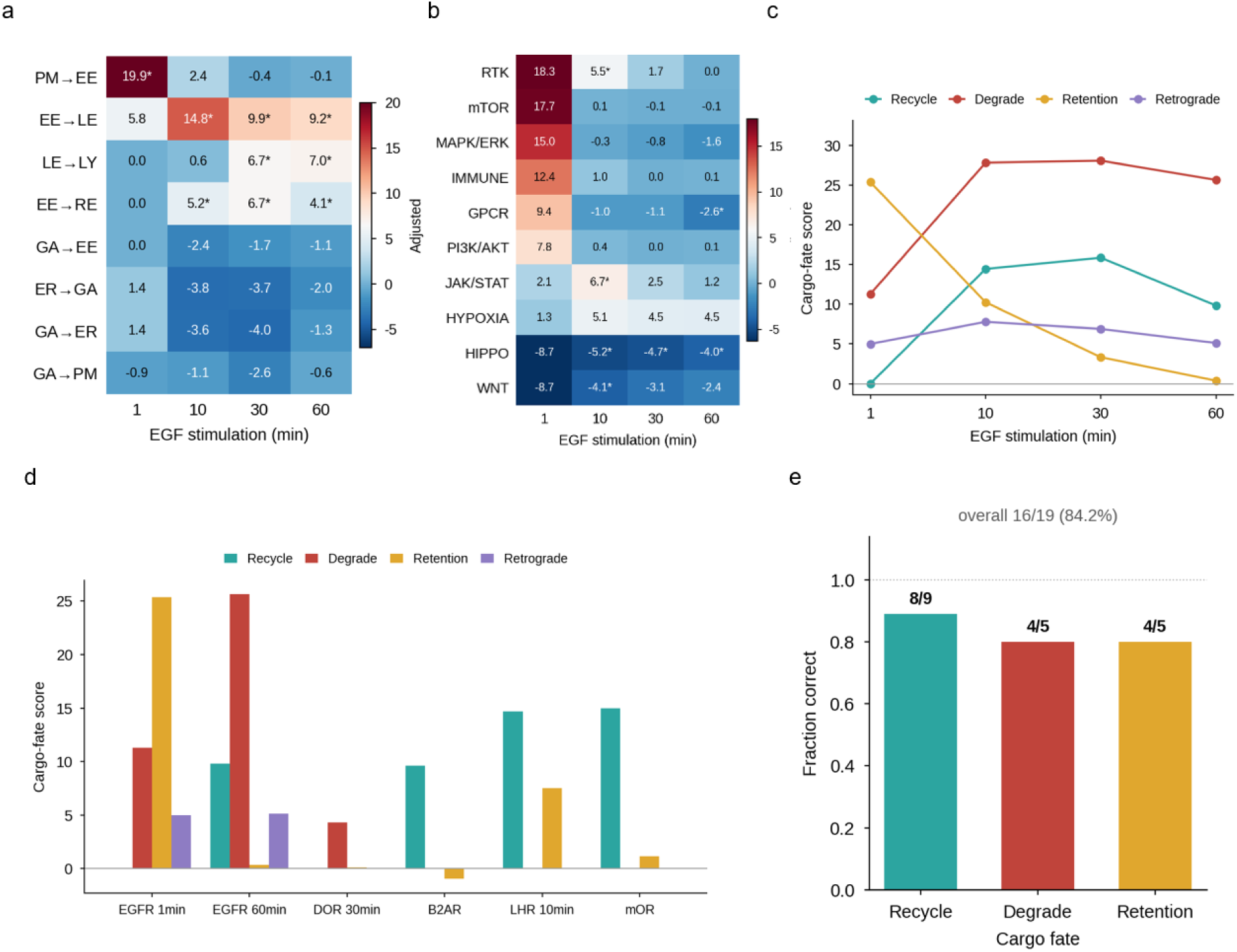
Directional scoring and cargo-fate inference. **a,b.** Transition and signaling-pathway scores across an EGFR-APEX2 time course at 1, 10, 30 and 60 min. Points are adjusted transition scores; asterisks indicate q < 0.05 after Benjamini–Hochberg correction within each dataset across the eleven transitions. **c.** EGFR cargo-fate scores across the same time course, projected onto the SORTER template. **d.** Fate-score profiles for six representative cargo conditions. **e.** Cargo-fate accuracy across 19 evaluable cargo-condition combinations spanning 11 cargoes: 8 of 9 recycling, 4 of 5 degradative and 4 of 5 retention conditions, 16 of 19 overall (84.2%). Expected fates were taken from the conclusions of the original publications before scoring.

The same transition scores were applied to score the same time course against the signaling reference set, comprising 1,837 proteins assigned to 22 pathways, revealing the highest receptor-proximal signaling-pathway scores at 1 min, followed by a decline towards baseline at 30–60 min (Fig. 3b). No individual pathway met the criteria for a specific pathway call, consistent with recruitment of signaling proteins shared across several pathways rather than selective engagement of a single pathway. Transition scores and signaling-pathway scores could therefore be evaluated from the same proteomic experiment. Projection of the transition scores onto the EGFR cargo template identified plasma-membrane retention as the highest-scoring fate at 1 min and degradation as the highest-scoring fate from 10 min onwards (Fig. 3c), consistent with ligand-induced EGFR downregulation [1]. A unique highest-scoring fate was obtained at each time point.

We next tested our cargo-fate inference methodology across 19 cargo-condition combinations representing 11 cargoes, and multiple published proximity-labeling and proteomic datasets [12,14–23]. For each condition, we assigned the expected fate based on the experimental design and published trafficking evidence. For the six representative cargoes shown in Fig. 3d, the highest-ranking supported fate matched the known behavior. The receptor conditions were evaluated in the context of stimulation or perturbation according to the original study; the transporter, channel, and resident-protein conditions were treated solely as transport-fate tests in and of themselves, not as ligand-activated or signal-regulated transporters or channels.

In the 19 conditions where the established fate was evaluable, we observed correct cargo-fate predictions in 16 (84.2% accuracy; Fig. 3e) including 8 of 9 recycling conditions, 4 of 5 degradation conditions and 4 of 5 retention conditions. The three incorrect predictions were the δ-opioid receptor at 10 min, where retention ranked above degradation, FGFR2b, where degradation ranked above recycling, and unstimulated GLUT4, where recycling ranked above retention. In each case, the expected fate was the next-highest supported fate.

Together, these results show that the transition profile inferred by Traffickome contains sufficient biological information to infer known membrane-cargo fates across a range of receptors, channels, transporters and compartment-resident-proteins. This analysis validates the route information captured by Traffickome, although it is not intended to replace direct imaging or biochemical measurements when the sole goal is to determine the transport of a single cargo.

### VPS35 perturbation provides a mechanistically interpretable case study

In the case of VPS35, we tested our method’s ability to predict loss of this core retromer subunit’s function, which is required for endosomal cargo recycling [3], as a mechanistically interpretable perturbation case study. Across a δ-opioid receptor proximity proteome [24] and an endosome-immunopurification proteome from iPSC-derived neurons [25], VPS35 removal consistently predicted its strongest effect on early endosome-to-recycling endosome transport (EE→RE; Fig. 4a).

**Figure 4.**
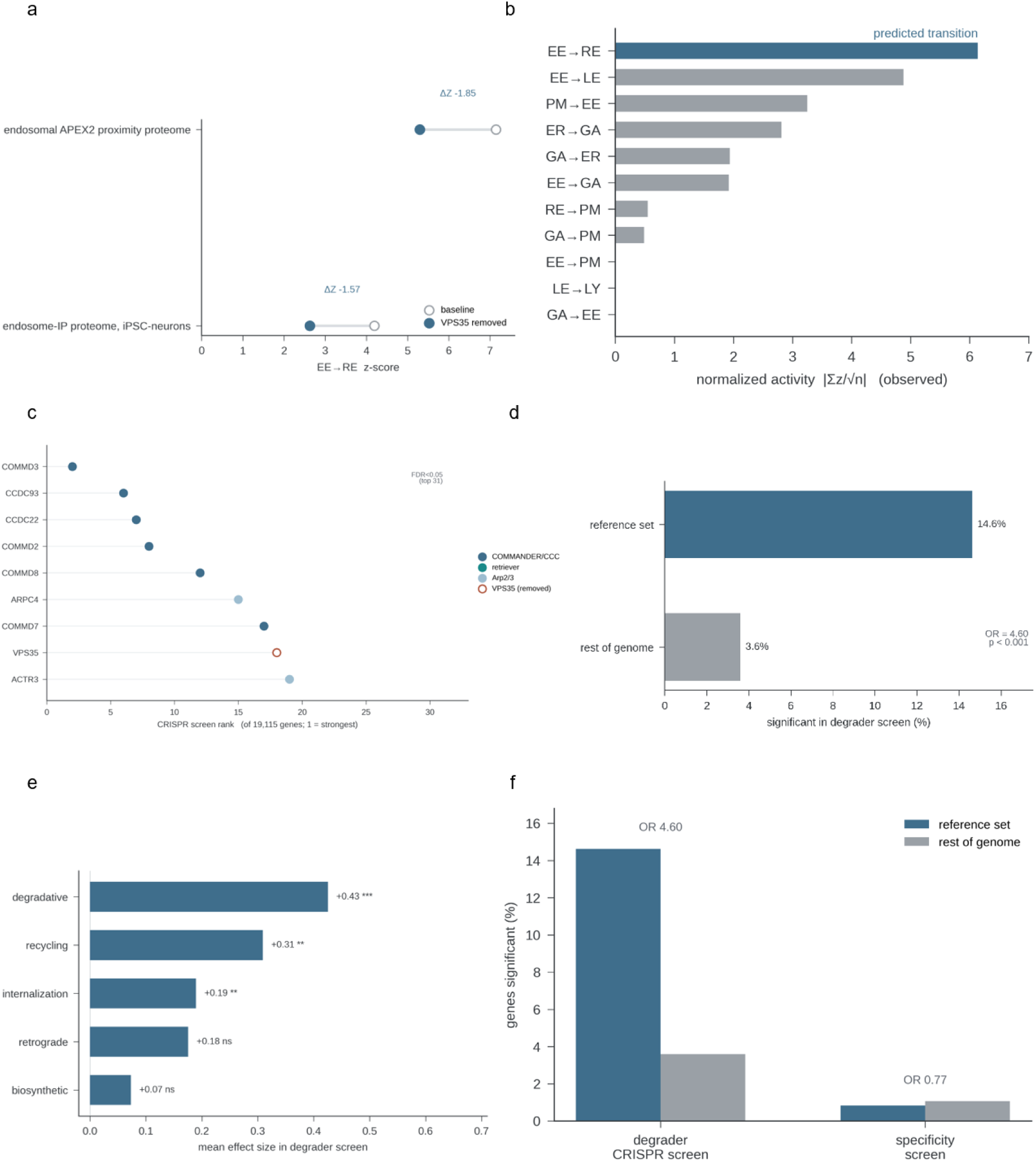
Perturbation case study and functional-genomics validation. **a.** Predicted EE→RE change after simulated VPS35 removal with propagation enabled, in a δ-opioid receptor proximity proteome and an endosome-immunopurification proteome from iPSC-derived neurons. **b.** All eleven transition scores in an experimental VPS35-knockout surface proteome; EE→RE ranks first (Pₑ = −6.138, q = 0.016). **c.** Machinery carrying residual EE→RE support after VPS35 removal, plotted against rank in an independent SARS-CoV-2 host-factor CRISPR screen. **d.** Reference-set genes among significant hits in a degrader CRISPR screen: 106 of 725 route-assigned genes (14.6%) versus 705 of 19,727 (3.6%); odds ratio (OR) 4.60, two-sided Fisher’s exact test, P < 0.001. **e.** Mean casTLE effect size in the degrader screen by cargo-fate class; ***P < 0.001, **P < 0.01, ns not significant, one-sample t-test against zero. **f.** The same comparison in a non-degradative specificity-control screen (OR 0.77, P = 0.71). OR, odds ratio; an OR above 1 indicates enrichment of reference-set genes among screen hits relative to the rest of the screened genome. Panels a–c assess mechanistic interpretability rather than independent validation of the curated VPS35 assignment.

We compared our predicted route-level change with a published cell-surface proteome from VPS35-knockout cells [26]. When the experimental dataset was analyzed using the standard transition-scoring procedure, EE→RE was the highest-ranked trafficking transition, with a normalized transition score of Pₑ = −6.138 (q = 0.016) (Fig. 4b). Thus, the experimental knockout showed the same route-level pattern as the simulation.

Propagating the simulated perturbation through the curated trafficking-complex hierarchy helped identify components of the CCC–Retriever/COMMANDER machinery, including the Retriever subunit VPS35L, together with Arp2/3-associated machinery, as contributors to the predicted recycling defect (Fig. 4c) [27]. These contributors also ranked among the strongest trafficking-associated hits in an independent host-factor CRISPR screen [28].

### Functional-genomics screens independently validate the trafficking reference set

For independent validation we analyzed two functional-genomics datasets that were not used to construct the trafficking reference set. In a genome-wide CRISPR screen for cellular factors required for lysosome-targeting-chimera-mediated degradation of membrane proteins [29], trafficking-reference proteins were more frequently identified as significant hits than non-reference genes, 106 of 725 route-assigned reference genes compared with 705 of 19,727 genes in the remaining screened background (14.6% versus 3.6%; odds ratio=4.60, two-sided Fisher’s exact test, P < 0.001; Fig. 4d).

This enrichment was concentrated among proteins assigned to degradation, recycling and internalization routes, whereas proteins assigned to the biosynthetic route were not enriched (Fig. 4e). This distribution was consistent with the lysosomal-degradation results of the screen. By contrast, a non-degradative specificity-control screen showed no enrichment of trafficking-reference proteins (odds ratio = 0.77; Fig. 4f). These results provide functional-genomic validation for both the composition and route-level organization of the Traffickome reference set.

### Traffickome distinguishes signaling associated with internalized and surface EGFR

Endocytosis determines where and for how long EGFR signals, but separating phosphorylation associated with internalized receptor from that associated with plasma-membrane receptor remains experimentally difficult because perturbations of endocytosis commonly affect additional cargoes and cellular processes. To distinguish these receptor pools, we used two experimental protocols to generate two tandem-mass-tag phosphoproteomic datasets in human oral squamous carcinoma HSC3 cells stimulated with 4 ng ml⁻¹ EGF.

In the first protocol, the cells were stimulated with EGF for 15 min, after which surface EGF–EGFR complexes were dissociated by a mild acid wash, thereby minimizing active EGFR in the plasma membrane and enriching active EGFR in endosomes (Fig. 5a).

**Figure 5.**
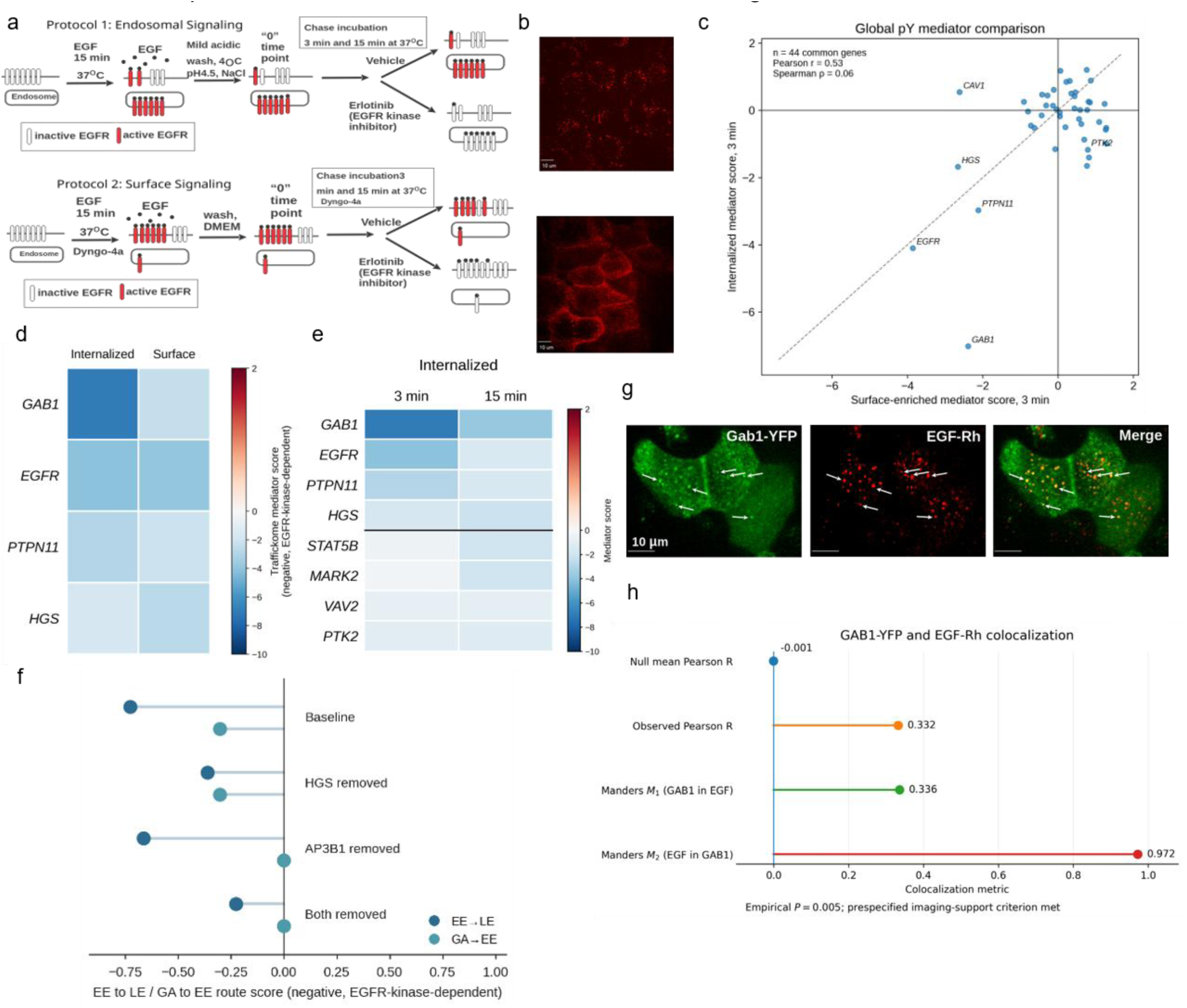
EGFR-dependent phosphorylation associated with internalized and surface receptor pools. **a.** Acid-wash and Dyngo-4a TMT 10-plex designs, with two independent measurements per condition. **b.** EGF-rhodamine localization confirming internalized and surface receptor pools; scale bars, 10 μm. **c–e.** Phosphotyrosine mediator scores compared between protocols and chase times; all four overlays were scored in fold-change-only mode to keep scoring identical across conditions. **f.** In silico perturbation in the 3-min internalized-receptor, phosphotyrosine-restricted overlay. Removal of HGS reduced EE→LE from −0.727 to −0.362 without altering GA→EE; removal of AP3B1 eliminated GA→EE with little effect on EE→LE; removal of both reduced EE→LE to −0.228. **g,h.** GAB1-YFP and EGF-rhodamine colocalization with Imaging-tab quantification: R = 0.332, null mean = −0.001, empirical P = 0.005 from 199 block randomizations, M1 = 0.336, M2 = 0.972. R, thresholded Pearson correlation; M1 and M2, thresholded Manders colocalization coefficients. One field, 8-bit channels; reported as exploratory evidence for overlap rather than as a compartment assignment.

In the second protocol, EGFR internalization was inhibited with Dyngo-4a,[30] enriching for phosphorylation associated with the receptor retained at the plasma membrane (Fig. 5a).

In both protocols, post-treatment of cells with Erlotinib, a small-molecule inhibitor of EGFR kinase, for 3 and 15 min was then used to identify phosphorylated proteins/sites that are dependent on EGFR kinase activity: sites maintained by active EGFR were expected to be dephosphorylated by protein tyrosine phosphatases (PTPs) following kinase inhibition.

Thus, these experiments were designed to distinguish phosphorylation maintained by internalized EGFR from phosphorylation maintained by plasma-membrane EGFR. Live-cell imaging of EGF-Rhodamine conjugate (EGF-Rh) confirmed predominantly endosomal localization following acid washing in the first protocol and increased plasma-membrane retention of active EGFR in Dyngo-4a-treated cells in the second protocol (Fig. 5b). It should be noted, however, that these protocols enriched for, but did not completely separate, surface- and endosome-associated receptor activity. Finally, each protocol used TMT10 peptide tagging and a ten-channel design comprising five chase conditions measured in two duplicate samples. The internalized-receptor protocol quantified 9,452 single-site phosphosite entries, whereas the surface-enriched protocol quantified 8,608 phosphosites across 2,227 proteins.

Figure 5c–e focuses on phosphotyrosine events, whereas Fig. 5f uses the 3-min internalized-receptor all-phosphosite overlay (Methods).

At the 3-min chase, GAB1 showed the largest negative mediator score in the internalized-receptor overlay (Fig. 5d). EGFR and PTPN11 were negative in both overlays, whereas HGS was more negative in the surface-enriched overlay. Thus, the two protocols produced overlapping but differently weighted phosphotyrosine profiles, consistent with enrichment rather than complete physical separation of the receptor pools. Across the 44 genes for which Traffickome produced a phosphotyrosine mediator score in both 3-min overlays, score magnitudes showed a moderate Pearson correlation but little rank-order agreement (Pearson r = 0.53; Spearman ρ = 0.06; Fig. 5c). This cross-protocol comparison is descriptive and does not imply that the same phosphosite was selected for a gene in both overlays.

The internalized-receptor phosphotyrosine profile also differed between chase times (Fig. 5e). GAB1, EGFR and PTPN11 showed stronger erlotinib sensitivity at 3 min, whereas HGS, STAT5B and MARK2 were more negative at 15 min; VAV2 and PTK2 showed smaller differences between the two chase times. These differences reflect site selection and dephosphorylation kinetics after EGFR kinase inhibition and should not be interpreted as sequential recruitment of “immediate” and “delayed” signaling proteins. Our published study independently established sustained EGFR–VAV2 signaling from endosomes,[31] providing mechanistic context for VAV2. In another example, Imaging of GAB1-YFP expressed in HeLa cells demonstrated strong colocalization of GAB1 with internalized EGF-Rh (Fig. 5g). Imaging-tab analysis of the same channels showed above-null colocalization in the analyzed field (thresholded Pearson R = 0.332, null mean = −0.001, empirical P = 0.005; M1 = 0.336, M2 = 0.972; Fig. 5h). Because the analysis used one 8-bit field and no compartment-marker channel, it was treated as exploratory evidence for GAB1–EGF overlap rather than as a compartment assignment or imaging–dataset concordance test.

At the transition level, the phosphotyrosine-restricted internalized-receptor analysis selected HGS pY289 as the representative phosphotyrosine event for HGS, which is a curated contributor to the early-endosome-to-late-endosome route (EE→LE), which directs activated EGFR towards degradation. For Fig. 5f, the 3-min internalized-receptor overlay was perturbed separately under phosphotyrosine restriction, giving a baseline EE→LE score of Pₑ = −0.727 and a baseline GA→EE score of −0.304. Simulated removal of HGS reduced EE→LE support from −0.727 to −0.362 without altering GA→EE, whereas removal of AP3B1 eliminated GA→EE support with little effect on EE→LE. Removing both reduced EE→LE further, to −0.228.

Although network propagation was enabled, no cascade term contributed to the reported EE→LE change. These results localized EGFR-kinase-dependent phosphorylation of HGS to the internalized receptor pool. The sorting function of HGS on this route has been well characterized. Our data indicate the compartment in which the associated phosphorylation occurs, thereby distinguishing signaling sustained after internalization from signaling at the plasma membrane.

### Traffickome identifies known trafficking transitions more accurately than conventional gene-set methods

To determine whether Traffickome ranks known trafficking transitions correctly, we analyzed 29 proteomic comparisons drawn from 14 published studies, spanning endocytic, recycling, degradative, secretory and retrograde transport.[12,14–18,26,32–38] For each comparison, we took the expected transition from the trafficking conclusion reported in the original publication rather than assigning it ourselves. The benchmark therefore tested whether Traffickome validates a published interpretation of the dataset. Because existing enrichment tools do not explicitly infer directed compartment-to-compartment transitions, we compared Traffickome with four methods implemented in decoupleR (ULM, MLM, ORA and GSEA), together with fgsea, single-sample GSEA, cameraPR and a simple over-representation baseline.[4–8] All methods received the same gene-level inputs and curated transition protein sets. The expanded benchmark added five endosomal proximity-proteome comparisons in which the bait protein was excluded from scoring: two VPS35 tag orientations, two VPS35L tag orientations and SNX27 [38]. For each included comparison, the expected transition was fixed as EE→RE before ranking. A broader seven-bait exploratory screen also contained SNX17 and HRS/HGS; their discordant Traffickome outcomes are reported in Supplementary Table 1.

Traffickome ranked the expected transition first in 23 of 29 comparisons (79.3%), compared with 17 of 29 comparisons (58.6%) for each of the three co-best enrichment methods (ULM, decoupleR GSEA and cameraPR) and 13 of 29 comparisons (44.8%) for the best over-representation baseline (Fig. 6a). For a paired inferential comparison, ULM was selected as a representative conventional enrichment model. Six discordant comparisons favored Traffickome and none favored ULM (exact two-sided McNemar test, P = 0.031). GSEA, cameraPR and the remaining methods are presented descriptively. Traffickome also showed consistent performance across transition classes represented by multiple datasets. The expected transition ranked first in all four endocytic datasets, 10 of 12 recycling datasets, all five degradative datasets and four of seven secretory datasets; none of the evaluated methods ranked the expected transition first for the single retrograde dataset, which is therefore omitted from Fig. 6b.

**Figure 6.**
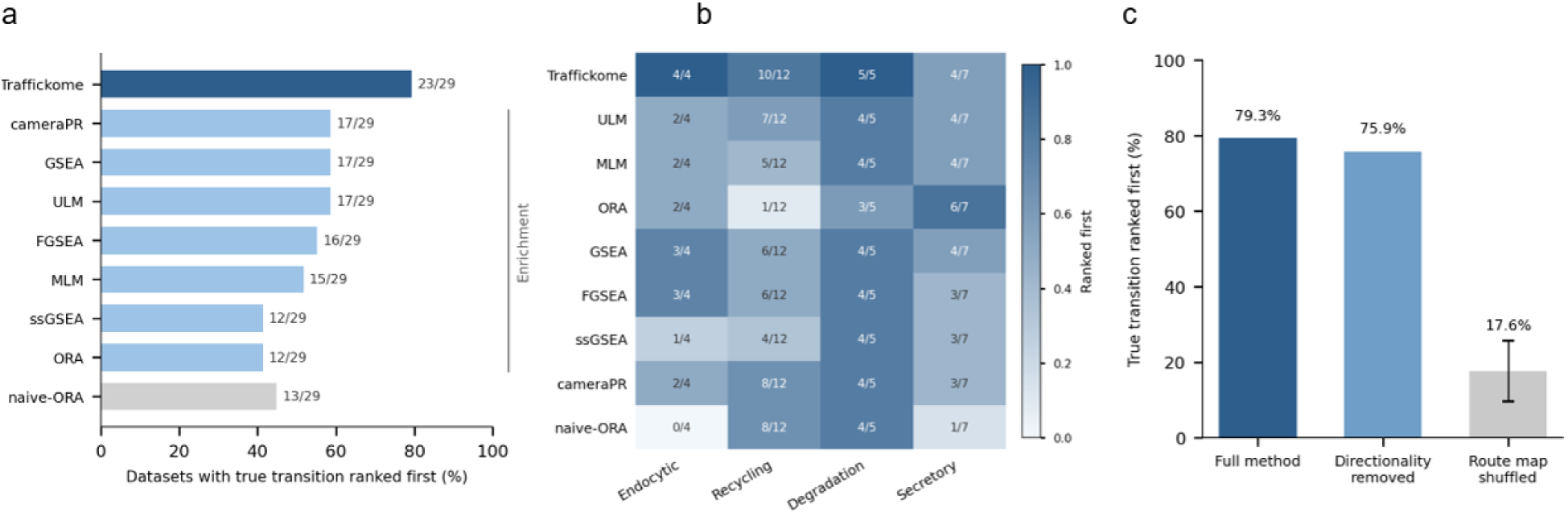
Benchmark of directed transition identification. **a.** First-rank accuracy across 29 proteomic comparisons from 14 published studies, with no-calls counted as incorrect so that all methods share the same denominator. Traffickome ranked the expected transition first in 23 of 29 comparisons (79.3%) versus 17 of 29 (58.6%) for the leading conventional gene-set methods; exact two-sided McNemar test P = 0.031 for Traffickome versus ULM, with six discordances favouring Traffickome and none favouring ULM. **b.** First-rank accuracy by transition class. The single retrograde comparison is not displayed because every method scored 0 of 1. **c.** Contribution analyses: replacing signed contributions with absolute values reduced accuracy to 75.9%, and randomizing protein-to-transition assignments while preserving transition sizes reduced it to 17.6% ± 8.1% across 20 randomizations (mean ± s.d.). ULM, univariate linear model.

Because every comparator received the same curated transition protein sets, we separated the contributions of the curated protein assignments and the direction of each protein’s change. Randomizing protein assignments among transitions while preserving transition sizes reduced mean correct identification from 79.3% to 17.6% ± 8.1% across 20 randomizations, indicating that the curated assignments accounted for most of the benchmark discrimination. Retaining the same protein assignments but scoring each protein’s change by magnitude alone, without its direction, reduced correct identification from 23 of 29 comparisons (79.3%) to 22 of 29 comparisons (75.9%). Direction is therefore encoded primarily in the curated assignment of proteins to directed transitions, which distinguish opposite routes between the same compartment pair, such as PM→EE from EE→PM. Whether each protein increased or decreased contributed one additional correct comparison (Fig. 6c).

## Discussion

Conventional enrichment analyses of proteomic data identify trafficking-associated proteins as a group but do not distinguish transport towards a compartment from transport away from it. In this study, we assigned trafficking proteins to defined compartment-to-compartment transitions and scored directional support from a single proteomic comparison, without spatial fractionation or model training. The framework contains three pairs of transitions that connect the same compartments in opposite directions, such as PM→EE and EE→PM. In 20 of the 29 comparisons the reverse of the correct transition was one of the available answers. Traffickome ranked the expected transition first in 79.3% of comparisons compared with 58.6% for the best conventional enrichment methods. Performance was weaker for secretory transport, where an over-representation baseline ranked the expected transition first in six of seven comparisons compared with four of seven for Traffickome. This is possibly due to secretory perturbations disrupting export machinery broadly rather than shifting a single directed step. The benchmark included one retrograde comparison and no method ranked the expected transition first, so directional inference on the secretory and retrograde routes will require datasets in which these routes are perturbed specifically. These results indicate that the direction of transport can be inferred from differential protein abundance for the endocytic, recycling and degradative routes.

Methods that infer differential localization from spatial proteomics ask whether a protein has changed compartment [9–11]. Directional inference asks which transport step between two compartments is engaged. Traffickome tells these apart because PM→EE and EE→PM are assigned different proteins. Consistent with this, randomizing which proteins belong to which transition reduced accuracy to 17.6%, whereas scoring each protein’s change by magnitude alone reduced it to 75.9%, indicating that direction is carried by the curated assignment of proteins to directed transitions rather than by the direction of individual protein changes. A practical consequence is that existing differential proteomic datasets can be interpreted for directional transport without new experiments, provided the relevant route is represented in the reference set. The reference set rather than the scoring procedure therefore limits where the method can be applied. Abundance and proximity-labeling experiments measure different biological quantities, proteins that were not measured cannot contribute to a score, and well-studied mammalian routes are better represented in the current reference set than context-specific mechanisms.

Cargo-fate projection translates transition scores into fate predictions that can be tested experimentally, and Traffickome classified 16 of 19 evaluable cargo-condition combinations correctly. In each case the expected fate ranked second. This is possibly due to overlapping cargo states, limited temporal resolution, or cargo-specific regulation that is absent from the current templates. Cargo-fate outputs are projections of the measured transition profile and do not measure cargo flux directly. Simulated loss of VPS35 predicted its strongest effect on early endosome-to-recycling endosome transport, and the same transition ranked highest when an experimental VPS35-knockout surface proteome was scored. Independent validation came from functional genomics, in which reference-set proteins were enriched among hits in a lysosome-targeting-chimera degradation screen, particularly among degradative, recycling and internalization assignments, and were not enriched in a non-degradative control screen.

Using Traffickome for compartment-enriched EGFR phosphoproteomics illustrates the combined interpretation of trafficking and signaling. The two receptor-pool enrichment protocols produced overlapping but differently weighted phosphotyrosine mediator profiles, and the internalized-receptor profile differed between chase times. These chase-time differences should be interpreted as site selection and dephosphorylation kinetics after EGFR inhibition rather than as a sequence of early and late recruitment events. In the phosphotyrosine-restricted analysis, HGS pY289 was the selected phosphotyrosine event for HGS within its protein-level EE→LE assignment. In a separate 3-min phosphotyrosine-restricted perturbation analysis, removal of HGS or of AP3B1 each reduced support for the transition to which that protein is assigned while leaving the other transition unchanged, and removal of both further reduced EE→LE score.

Because EGFR is a tyrosine kinase, the serine and threonine sites in that analysis reflect downstream kinase activity rather than direct EGFR phosphorylation. These findings associate continued EGFR kinase activity with phosphorylation changes in receptor-sorting machinery. However, acid washing and Dyngo-4a enriched for internalized and surface receptors without separating the two pools completely. Erlotinib-sensitive phosphotyrosine changes indicate dependence on EGFR kinase activity but do not identify direct EGFR substrates. The mediator comparisons were based on fold change alone, because per-site P values derived from replicates were not available for all four overlays.

## Methods

### Directional reference framework

Traffickome uses a curated reference set of 754 trafficking proteins, 752 of which are assigned to one or more of 11 directed transitions across seven route-endpoint compartments, together with 1,837 signaling proteins assigned to 22 pathways. Protein-to-transition assignments were derived from Gene Ontology, Reactome, UniProt and manual review of primary literature.[39–41] The signaling reference set additionally draws on KEGG and MSigDB,[42,43] with HGNC used for gene-symbol normalization.[44] Directional phosphosite rules were curated separately using primary literature, Reactome trafficking context and PhosphoSitePlus regulatory-site annotations; PhosphoSitePlus was not used to define protein-to-transition membership.[40,45] Thirty-nine trafficking complexes and 20 directed complex dependencies support the perturbation analysis; a 420-protein trafficking-machinery subset is used for phosphoproteomic transition scoring. Protein assignments, directional phosphosite rules, complex membership and source identifiers are provided in the deposited reference files.

### Input processing and transition scoring

Uploaded tables are classified as signed differential, fold-change-only, paired-intensity, single-intensity or unsigned hit-list inputs. Gene symbols are resolved through the bundled HGNC alias table. Event-level evidence is derived from the available fold change, P value or intensity measurements; when multiple events map to one gene, the event with the largest absolute score is retained while preserving its sign. Gene scores are standardized within each dataset. For phosphosites with a directional rule, the site score is oriented according to whether phosphorylation promotes or inhibits the specified transition.

For transition e, the canonical score is Tₑ = Σz_g/√nₑ, where z_g is the standardized score of each measured transition contributor and nₑ is the number of measured contributors. Transition scores are then mean-centred across the 11 transitions and multiplied by a coherence term, defined as the absolute sum of contributor z scores divided by the sum of their absolute values. This adjusted score is the statistic used for ranking and for significance testing. Contributor-count-matched permutation nulls (1,000 permutations) provide normalized scores and empirical P values. Benjamini–Hochberg correction is applied within each dataset across the 11 transitions, which are not independent because of the mean-centring step.[46] SPECIFIC, BROAD and NONE classifications are based on the adjusted P value. Full input-mode equations, contributor definitions, ranked-output rules and robustness calculations are provided in Supplementary Method 2.

### Cargo-fate and perturbation analyses

Cargo-fate projection maps normalized transition scores onto cargo-specific routing templates with allowed branches for recycling, degradation, retention or retrograde transport. Fate scores are standardized with a within-template permutation null, and a no-call is returned when the template lacks measured support or the leading fate is not unique. Cargo-fate outputs are projections of transition-level evidence rather than direct measurements of cargo flux.

The perturbation analysis retains gene-level event magnitudes, modifies the selected contributor or contributor set, and recalculates affected transition scores against contributor-count-specific nulls. Knockout simulations set the direct contribution to zero. When enabled, losses are propagated once through the curated complex-dependency hierarchy and applied only to measured downstream contributors. Multi-gene perturbation applies the intervention simultaneously to the requested genes. Complete equations and eligibility rules are provided in Supplementary Method 4.

VPS35, HGS and AP3B1 perturbations were used as mechanistically interpretable case studies rather than independent validation of assignments that contributed to construction of the curated framework.

### Image colocalization analysis

For the GAB1-YFP/EGF-rhodamine analysis, the two channels were thresholded with a Costes-inspired paired regression-line procedure.[47] Pearson correlation and thresholded Manders coefficients[48] were calculated over pixels exceeding both thresholds. Spatial significance was evaluated by 199 10-pixel block randomizations. The single analyzed field contained no compartment-marker channel; it therefore supported GAB1–EGF colocalization but was not used to assign a compartment, validate a transition endpoint or calculate imaging–dataset concordance. Extended imaging functions are described in Supplementary Method 5.

### Benchmark and validation datasets

Transition identification was evaluated using 29 proteomic comparisons from 14 published studies with expected trafficking classes and acceptable directed transitions specified before scoring. A comparison was correct when the highest-ranked absolute canonical transition score belonged to the acceptable set. Dataset identities, source publications, expected transitions and predicted transitions are listed in Supplementary Table 1, and cargo-fate conditions in Supplementary Table 2.

Traffickome was compared with ULM, MLM, ORA and GSEA through decoupleR, together with fgsea, single-sample GSEA and cameraPR,[4–8] using the same gene-level inputs and transition sets. Controls replaced signed contributions by their absolute values, randomized protein-to-transition assignments while preserving transition sizes, and removed 10–50% of reference proteins to assess incomplete annotation. The paired Traffickome–ULM comparison used an exact two-sided McNemar test; other method comparisons were descriptive.

Cargo-fate validation used 19 prespecified cargo-condition pairs spanning 11 cargoes. Perturbation case studies used VPS35-associated endosomal datasets and the 3-min internalized-receptor phosphoproteomic overlay. Functional-genomics corroboration used a lysosome-targeting-chimera CRISPR screen and DepMap gene-effect profiles.[29,49] Detailed dataset processing and statistical procedures are provided in Supplementary Methods 6 and 7.

### Compartment-resolved EGFR phosphoproteomics

HSC3 cells were grown in DMEM containing 5% FBS and were serum-starved before stimulation. In the internalized-EGFR-enriched protocol #1, confluent cells in 15-cm dishes were stimulated with EGF at 4 ng ml⁻¹ for 15 min at 37 °C, and EGF-EGFR complexes remaining at the cell surface were dissociated by a 2-min incubation in ice-cold 0.2 M acetic acid (pH 4.5).

Cells were rinsed with ice-cold DMEM to neutralize the acid wash and were either solubilized immediately or chased with Erlotinib (0.5 μM) or DMSO (vehicle) for 3 or 15 min at 37 °C before lysis. In the surface-EGFR-enriched protocol, EGFR internalization was inhibited by preincubation with 30 μM Dyngo-4a,[30] and the cells were stimulated with 4 ng ml^−1^ EGF for 15 min at 37^°^C in the continued presence of Dyngo-4a followed by Erlotinib or vehicle chase as in protocol #1. Finally, cells were lysed in the TGH buffer as described previously,[31] and the resulting 10 lysates from each protocol were cleared by centrifugation at 16,000g. Proteins were digested by trypsin and LysC, and phosphopeptides were enriched using a titanium bead-based column as described [50]. Quantitation was performed using TMT 10-plex and fractionated samples were analyzed on an Orbitrap Fusion as described [50]. Each protocol yielded ten TMT channels: two independent measurements for vehicle and erlotinib chases at 3 and 15 min, plus the corresponding baseline/control channels according to the experimental design.

For each protocol and chase time, the Traffickome input was the site-level log₂ ratio of erlotinib to vehicle. Peptides containing more than one annotated residue were expanded into separate site records. When multiple sites mapped to one gene, the event with the largest absolute event score was retained during gene-level collapse, preserving its sign.

For the receptor-proximal mediator analyses in Fig. 5c–e, inputs were restricted to phosphotyrosine sites before gene-level collapse. To maintain identical scoring across protocols and chase times, all four phosphotyrosine overlays were analyzed in fold-change-only mode; available P values from the surface-enriched protocol were not used in the primary comparison. The canonical EE→LE assignment was likewise phosphotyrosine restricted. The route perturbation in Fig. 5f was calculated separately from the 3-min internalized-receptor all-phosphosite overlay, which included serine, threonine and tyrosine sites.

A negative mediator or perturbation score indicated decreased phosphorylation after erlotinib and therefore dependence on continued EGFR kinase activity in the corresponding compartment-enrichment condition. Erlotinib sensitivity does not establish direct phosphorylation by EGFR: serine/threonine events are necessarily indirect, and phosphotyrosine events may reflect downstream kinases or phosphatase-dependent persistence. The experiment contained two independent measurements per condition and was interpreted primarily from effect magnitude, consistency between measurements, transition rank and agreement across chase times and compartment-enrichment protocols.

For Fig. 5h, GAB1-YFP was analyzed as the protein-of-interest channel and EGF-rhodamine as cargo in one 8-bit field. Thresholded Pearson correlation, thresholded Manders coefficients and an empirical P value from 199 deterministic block randomizations were reported. No compartment-marker channel was supplied; therefore the analysis supported GAB1–EGF colocalization but did not assign a compartment, validate a transition endpoint or calculate imaging–dataset concordance.

### Implementation, statistics and reproducibility

The browser and Python implementations use synchronized versioned reference resources. Transition, pathway, cargo and perturbation calculations are deterministic for fixed inputs, parameters and reference version. Transition and pathway empirical P values use 1,000 permutations unless otherwise stated; imaging uses 199 block randomizations and contributor-dropout robustness uses 500 iterations. Anthropic Claude and OpenAI GPT models were used as coding assistants during development of the Traffickome framework. All model-assisted code was reviewed, tested and validated by the authors, who take responsibility for the software and analyses.

Unless otherwise stated, statistical tests were two-sided. Benjamini–Hochberg correction was applied within each dataset. Benchmark performance used the common comparison denominator, with no-calls counted as incorrect. No formal power calculation determined the number of public benchmark datasets; inclusion was constrained by interpretable trafficking ground truth and reusable quantitative data.

Dataset-specific preprocessing, thresholds, benchmark labels and analysis workflows are reported in Supplementary Methods, Supplementary Tables 1 and 2, and the figshare deposit.

### Declaration of generative AI and AI-assisted technologies in the manuscript preparation process

During the preparation of this manuscript the authors used ChatGPT (OpenAI) and Claude (Anthropic) in order to improve language and readability, to assist with typesetting and formatting, and to assist with code development for the Traffickome computational framework. After using these tools the authors reviewed and edited the content as needed. All data were generated by the authors, all analyses were designed and executed by the authors, and all reported values were verified against source data by the authors, who take full responsibility for the content of the published article.

## Data and code availability

The Traffickome browser application, Python package, source code and versioned reference files are deposited at figshare and will be publicly released on publication; the DOI will be provided once issued. The deposit contains the curated reference sets, the published proteomic comparisons analyzed here, the transition and cargo-fate ground-truth files, and both implementations of the software. Public datasets are redistributed where permitted, and their original accessions and publications are given in the cited source references.

The compartment-resolved EGFR phosphoproteomic data will be deposited through ProteomeXchange/PRIDE, and the processed overlays used as Traffickome inputs will be added to the figshare deposit, on publication. Processed phosphosite tables used as Traffickome inputs will be provided as Source Data. The CRISPR-screen and DepMap data are available from their original sources [29,49].

## Funding

This work was supported by National Institute of General Medical Sciences of the National Institutes of Health grant R35GM148363 to Alexander Sorkin.

## Author contributions

R.D.: Conceptualization, data curation, formal analysis, investigation, methodology, software, validation, visualization and writing (original draft, review and editing). I.P.-M.: Investigation, methodology, resources and writing (review and editing). J.P.: Investigation, methodology and writing (review and editing). S.P.G.: Resources, supervision and writing (review and editing). A.S.: Conceptualization, funding acquisition, project administration, resources, supervision, visualization and writing (review and editing).

## Competing interests

The authors declare no competing interests.

## Notes

### Competing Interest Statement

The authors have declared no competing interest.

