## Supplementary Methods for "Traffickome infers directed, compartment-resolved membrane trafficking from proteomic data"

### Supplementary Method 1. Reference sets and curation

The trafficking reference set contains 754 proteins, of which 752 are assigned to at least one of eleven directed transitions: PM→EE, EE→PM, EE→RE, RE→PM, EE→LE, LE→LY, EE→GA, GA→EE, ER→GA, GA→ER and GA→PM. These transitions span seven of ten defined compartments; the remaining three are used for annotation only and are not route endpoints. UniProt accessions identify each protein.

Transition membership was assigned from Gene Ontology Biological Process terms and Reactome pathway annotations, and compartment membership from Gene Ontology Cellular Component terms.189 assignments, covering 174 proteins, were refined by manual review of the primary literature. All contributors to a transition are scored equally.

The signaling reference set contains 1,837 proteins assigned to 22 named pathways. 16 canonical signaling pathways were assembled from KEGG signal-transduction pathways and the Reactome Signal Transduction hierarchy, and pathway membership was filtered against UniProt functional annotations. Six further gene sets covering cancer and stress-response programs were taken from the MSigDB Hallmark collection.

39 multisubunit trafficking complexes and twenty directed dependencies between them support the perturbation analysis, with complex membership curated manually against the primary literature. A 420-protein machinery subset is used as the permutation pool for phosphoproteomic transition scoring.130 directional phosphosite rules were curated from primary literature, Reactome trafficking context and PhosphoSitePlus regulatory-site annotations, each retaining its supporting PubMed identifiers. PhosphoSitePlus was not used to define protein-to-transition membership. Gene symbols were resolved through the HGNC complete set.

### Supplementary Method 2. Input processing and transition scoring

The input is a table of gene identifiers with, depending on the input mode, log2 fold changes, P values, one or two intensity measurements, or an unsigned gene list. Phosphoproteomic inputs may additionally carry residue positions. The importer resolves gene symbols through the bundled HGNC alias table, detects the data modality, and assigns one of five input modes: signed differential, fold-change only, paired intensity, single intensity, or hit list.

A gene contributes when at least one of its events satisfies the selected gates. Default gates are an absolute log2 fold change of at least 0.5 and a P value of 0.05 or below; in fold-change-only and hit-list modes the P-value gate is not applied. When several events map to the same gene, the event with the largest absolute score is retained and its sign preserved. Gene scores are standardized within each dataset:

$$z_\gamma = (s_\gamma - \mu_s) / \sigma_s \qquad (1)$$

where $s_\gamma$ is the retained score for gene g, and $\mu_s$ and $\sigma_s$ are the mean and population standard deviation across all genes with defined scores. If $\sigma_s$ is zero, all standardized scores are set to zero and a warning is issued. For phosphosites carrying a directional rule, the sign of $z_\gamma$ is oriented according to whether phosphorylation promotes or inhibits the transition.

For transition e, the canonical transition score is

$$T_e = \Sigma z_\gamma / \sqrt{n_e} \quad (2)$$

where the sum runs over the measured contributors assigned to that transition and $n_e$ is their number. For phosphoproteomic inputs, only contributors within the trafficking-machinery subset are included. Coherence measures sign agreement among contributors:

$$\kappa_e = |\Sigma z_\gamma| / \Sigma |z_\gamma| \quad (3)$$

Canonical scores are then centred on their mean across all eleven transitions, with transitions having no measured contributor entering the mean as zero, and multiplied by coherence:

$$A_e = (T_e - \bar{T}) \times \kappa_e \quad (4)$$

Equation (4) is the statistic used for ranking, significance testing and classification.

Significance is assessed by permutation. In each of 1,000 permutations, the eligible gene scores of the dataset are randomly shuffled and partitioned into consecutive blocks matching the contributor count of each transition, and equations (2) to (4) are recomputed on the permuted blocks. The same shuffle is applied to all eleven transitions within a permutation, so the null preserves both the contributor counts and the total pool. For phosphoproteomic inputs the pool is restricted to the trafficking-machinery subset. Two-sided empirical P values are the number of permutations in which the absolute permuted statistic meets or exceeds the observed value, plus one, divided by 1,001. Benjamini–Hochberg correction is applied within each dataset across the scored transitions, which are not independent because of the mean-centring in equation (4), and adjusted values are enforced to be monotone in rank and capped at one.

Transitions are ranked by the absolute adjusted score, with the sign of the summed contributor scores giving the direction. A transition is classified SPECIFIC when its adjusted P value is below 0.05, BROAD when it is scored but does not meet that threshold, and NONE when no contributor is measured or the adjusted score is zero. The ranked panel reports these values together with the highest-contributing genes and applies no further test.

Robustness to incomplete measurement is assessed by randomly removing 30% of the unique measured contributors in each of 500 iterations, applying the same retained-gene set to all eleven transitions and rescoring with equation (2). The diagnostic reports the median resampled score, a percentile interval, direction stability, rank stability and top-k retention. It generates no new P values and does not alter the reported classifications.

**Supplementary Method 3. Cargo-fate projection**

Each cargo is assigned a template that defines the allowed fates for that cargo and, for each fate, the directed transitions composing its branch together with a weight. Cargo names are resolved to templates through a curated cargo table, which carries aliases and gene-symbol equivalents so that receptor shorthand resolves to the corresponding gene.

For the template branch representing fate f, the fate score is

$$F_e = \Sigma\, w_t\, A_t \quad (5)$$

where the sum runs over the transitions in that branch, $w_t$ is the curated weight of transition t within the branch, and $A_t$ is the adjusted transition score from equation (4). No cargo-specific multipliers, coupling penalties or additional size normalization are applied, since the template structure specifies both the transitions and their relative contribution.

The predicted fate is the allowed fate with the largest score. The margin is the difference between the largest and second-largest fate scores and indicates how clearly the leading fate is separated. A cargo-condition case is evaluable only when at least one transition in the template has measured support and one allowed fate has a uniquely largest score; cases with no measured support in the template, or with tied maxima, are returned as no-calls.

Cargo-fate projection was evaluated on 19 cargo-condition combinations spanning 11 cargoes, of which 16 were classified correctly, with the established fate assigned before scoring from the conclusions of the original publications. Accuracy was calculated among evaluable cases. Quantitative datasets were projected in their native input mode, and the two unsigned hit lists were evaluated separately, with listed genes treated as equal positive evidence.

Cargo-fate outputs are projections of the measured transition profile onto curated routing templates. They do not measure cargo flux and are not intended to replace imaging or biochemical trafficking assays for individual cargoes.

**Supplementary Method 4. In silico perturbation**

The perturbation analysis estimates the effect of removing a trafficking protein on the transitions to which it contributes. Knockout removes the selected gene from the contributor sum of every transition it belongs to. Only transitions containing the selected gene, or a gene affected by propagation, are recomputed.

Unlike equation (2), the perturbation statistic normalizes by the complex-adjusted contributor count n_eff, in which contributors belonging to the same curated complex together contribute one unit while subunits shared between complexes are counted individually:

$$P_e = \Sigma\, z_\gamma \,/\, \sqrt{n\_eff} \quad (6)$$

The perturbed score is standardized against a contributor-count-matched null, giving

$$Z_e = (|P_e| - \mu_n) \,/\, \sigma_n \quad (7)$$

where $\mu_n$ and $\sigma_n$ are the mean and standard deviation of the null for the corresponding contributor count, and $Z_e$ is capped at a fixed maximum to prevent unstable values when the null distribution is narrow. Results are reported as the change in $P_e$ and the change in $Z_e$ relative to baseline, ordered by the absolute change in $Z_e$. Transitions whose absolute changes in both quantities fall below 0.001 are omitted. A direction change is reported when the baseline and perturbed scores are both nonzero and of opposite sign.

Complex-dependency propagation models the effect of losing one subunit on functionally dependent complexes. For a gene belonging to a complex of N members, knockout imposes an initial functional loss of 1/N on that complex. For a directed dependency from complex c to complex d with curated strength a, the transmitted loss is the product of the loss at c and a, and losses arriving at the same downstream complex combine as independent contributions so that the combined loss is one minus the product of their complements. Losses are applied in a single pass in topological order. Each measured member of an affected complex then receives a multiplier equal to one minus the loss of that complex, and where a gene belongs to several affected complexes the lowest multiplier is retained. The perturbed gene itself is excluded from this calculation. Only propagated genes that are measured and eligible in the selected dataset contribute to the recalculated scores. Propagation is enabled by default and can be disabled to obtain the direct effect of removing a single gene.

A multi-gene perturbation applies the same intervention to a defined set of genes, such as the measured members of one complex. A separate cascade is computed for each member and the resulting multipliers are combined by retaining the strongest predicted reduction. At least one requested gene must contribute to the baseline under the selected dataset and gates; otherwise the perturbation is returned as a no-call rather than as a zero effect.

**Supplementary Method 5. Image colocalization**

Colocalization was calculated from a single two-channel fluorescence field. Intensity thresholds for the two channels were set by Costes regression sweep, in which candidate threshold pairs are tested from the maximum intensity downward and the first pair is retained for which the Pearson correlation of pixels below both thresholds is not positive. The analysis mask comprises pixels above either threshold. Thresholded Pearson correlation and thresholded Manders coefficients M1 and M2 were computed over the masked pixels.

Spatial significance was assessed by 199 Costes block randomizations, in which one channel is divided into 10-pixel square blocks that are randomly rearranged and the thresholded Pearson correlation is recalculated. The empirical P value is the proportion of randomizations reaching or exceeding the observed correlation. Because a single field was analysed, the result is reported as endpoint localization evidence and not as a measure of transport direction or kinetics.

**Supplementary Method 6. Benchmark and validation datasets**

Transition identification was evaluated using 29 proteomic comparisons from 14 published studies. For each comparison, an expected trafficking class and one or more acceptable directed transitions were assigned before scoring, taken from the trafficking conclusions of the original publications. The benchmark comprised four endocytic, twelve recycling, five degradative, seven secretory and one retrograde comparison. Dataset identities, source publications, conditions, input modes, acceptable transitions and predicted transitions are listed in Supplementary Table 1; cargo-fate conditions and predictions are listed in Supplementary Table 2.

Each dataset was scored in its native input mode, and the highest-ranked transition was compared with the acceptable set. A method returning no rankable transition for a dataset was counted as incorrect, so all methods were evaluated on the same 29-comparison denominator. Traffickome was compared with univariate and multivariate linear models, over-representation analysis, gene-set enrichment analysis, fgsea, ssGSEA and cameraPR, each supplied with the same transition protein sets, and with a naive over-representation baseline. Performance was summarized overall and by transition class. Paired method comparisons used the exact two-sided McNemar test on discordant comparisons.

Two reanalyses separated the contribution of the reference set from that of the scoring procedure. In the first, protein-to-transition assignments were randomized while preserving the number of proteins per transition, across 20 randomizations. In the second, the sign of each gene-level change was excluded and contributors were scored by magnitude alone, retaining the same assignments.

Enrichment of reference-set proteins among functional-genomics hits was evaluated by two-sided Fisher's exact test against the remaining screened genes, using the 752 proteins carrying a directed-transition assignment, and was repeated on a non-degradative screen as a specificity control.

**Supplementary Method 7. Statistics, determinism and availability**

Statistical tests were two-sided unless stated otherwise. Transition, pathway, cargo-fate and perturbation null distributions used 1,000 randomizations. Benjamini–Hochberg correction was applied within each dataset. No formal procedure was used to determine the number of public benchmark datasets; all datasets meeting the eligibility criteria were included. The compartment-resolved EGFR phosphoproteomic experiment contained two independent measurements per condition and was interpreted from effect magnitude, consistency between measurements and transition rank rather than from per-site significance.

Traffickome was implemented as a browser-native JavaScript application and as an installable Python package, both using the same versioned reference files, cargo templates, compartment definitions, phosphosite rules, HGNC alias table and complex hierarchy. The browser performs all scoring locally and uploaded data are not transmitted to a remote server. Transition, pathway, cargo-fate and perturbation calculations are deterministic for fixed input data, input mode, gates and reference-set version.
